# Analysis of *lhx8a, isl1, pax6a/b, calb2a* and *sst7* Reveals that Dopaminergic Neurons in the Zebrafish Subpallium Belong to the Extended Amygdala

**DOI:** 10.1101/2025.01.27.635026

**Authors:** Daniel Armbruster, Thomas Mueller, Wolfgang Driever

## Abstract

The amygdala is a heterogenous multinuclear telencephalic structure critical for motivated and emotion-related behaviors in vertebrates. In ray-finned fish (actinopterygii) like the teleost zebrafish, a telencephalic outward growing process called eversion makes defining amygdaloid territories particularly challenging. Teleosts are also peculiar in that they develop prominent dopaminergic neuron groups in the subpallium, which are absent from tetrapods. To shed light on amygdala organization in teleosts, we pursued an evolutionary-developmental approach focusing on the topological origin of subpallial dopaminergic neurons. Specifically, we analyzed developmental expression patterns of Tyrosine hydroxylase in conjunction with *pax6a+b*, *isl1a*, *nkx2.1*, *lhx8a, calb2a* as telencephalic topology markers in brains of 5- and 30-day-old zebrafish (*Danio rerio*, Teleostei). Our results reveal, that the subpallial dopaminergic neurons develop within a *pax6a* negative dorsal subpallial domain (Vdd), which forms a primordial portion of the extended amygdala, including the medial amygdala and the anterior bed nucleus of the stria terminalis. Moreover, these dopaminergic neurons differentially coexpress *calb2a* and *sst7*, indicating population heterogeneity and potentially functional diversity. Our data also show that the zebrafish extended amygdala is formed by the dorsal LGE-like Vdd, which is subdivided into a Vdd2 subdivision that may correspond to the extended medial amygdala, including the bed nucleus of the stria terminalis, and forms the pallial-subpallial border region, and a more ventral Vdd1 that is *pax6a* positive and that corresponds to the central amygdala. Our work contributes to understanding development and evolution of the amygdala, and provides a foundation for functional analysis of the newly defined dopaminergic subtypes of the extended amygdala.

## 3. Introduction

Dopaminergic (DA) neurons establish several highly conserved neuromodulatory systems in the vertebrate brain (Smeets & Gonzalez, 2000; Yamamoto & Vernier, 2011). Highly conserved from fish to mammals are DA systems in the retina, olfactory bulb, hypothalamus and the ventral diencephalon. In addition, amphibs, birds and mammals have prominent mesdiencephalic DA systems projecting to cortical and limbic structures, where they are involved in the regulation of learning and memory processes, reward system and fear responses, as well as to the striatum, contributing to movement control pathways of the basal ganglia (Bjorklund & Dunnett, 2007; Iversen & Iversen, 2007; Bromberg-Martin et al., 2010) Strikingly, these substantial populations of mesdiencephalic DA neurons are not conserved in all vertebrate taxa, since they are not present in lamprey and ray-finned fish (Actinopterygii) (Pierre et al., 1997). For the widely used teleost zebrafish model, the absence of functional midbrain DA neurons has been well documented (Holzschuh et al., 2001; Kaslin & Panula, 2001; Filippi et al., 2010; Yamamoto et al., 2010). However, recently, sparse cells that weakly express Tyrosine hydroxylase (Th), a marker for catecholaminergic neurons, have been detected in the midbrain of various teleosts (Lopez et al., 2019; Lozano et al., 2019; Borgonovo et al., 2021; Altbürger et al., 2023). In zebrafish, it was shown that they do not express other markers required for DA signaling (Altbürger et al., 2023), indicating that these midbrain cells may be non-functional with regard to DA neuromodulation. However, unlike tetrapods, teleosts have a well-developed group of DA neurons in the subpallium (Guo et al., 1999; Holzschuh et al., 2001; Rink & Wullimann, 2002; Filippi et al., 2010; Yamamoto & Vernier, 2011). Interestingly, also in the striatum of rodents and primates, including humans, Th-immunoreactive neurons have been identified that were classified as GABAergic Th-interneurons (THINs) (Dubach et al., 1987; Tashiro et al., 1989; Porritt et al., 2000; Ibanez-Sandoval et al., 2010). However, further characterization of THINs in rodents revealed that they neither express other proteins required for functional DA neuromodulation, nor they release DA upon optogenetic activation (Xenias et al., 2015). The presence of dysfunctional THINs and the mesdiencephalic systems in mammals compared to the subpallial DA system and dysfunctional midbrain DA neurons in zebrafish suggests a divergent evolution of DA systems in vertebrates. Several species of cartilaginous fish (Chondrichthyes) have DA neurons in areas corresponding to ventral tegmental area and substantia nigra (Stuesse et al., 1994; Meredith et al., 1999), suggesting that teleosts may have secondarily lost midbrain DA neurons.

This raises the question about potential sources of DA in the zebrafish pallium and subpallium. Apart from a few projections of a small subset of the diencephalic DA cells in the posterior tuberculum, no extratelencephalic DA input into the zebrafish telencephalon proper has been shown (Rink & Wullimann, 2001; Tay et al., 2011). Another strong candidate for telencephalic DA modulation in zebrafish are the subpallial DA neurons, since they were shown to form extensive local intratelencephalic arborizations (Tay et al., 2011). Like THINs of mammals, the zebrafish subpallial DA neurons have a GABAergic co-transmitter type, consistent with recent findings in goldfish (Filippi et al., 2014; Tibi et al., 2023). Unfortunately, so far there is limited knowledge of anatomical localization, function and connectivity of these subpallial DA cell populations. Due to their topographical localization in the dorsal and central subpallium (Vd, Vc), these cells have been discussed as being in the zebrafish equivalent of the striatum (Tay et al., 2011). Recently, however, we proposed that the subpallial DA neurons of zebrafish are located in a region of the teleostean equivalent of the subpallial extended amygdala (EA) we specified as the anterior bed nucleus of the stria terminalis (BSTa) (Porter & Mueller, 2020).

The amygdala has mostly been investigated in tetrapods, which revealed that it is composed of a number of pallial and subpallial nuclei with interspersed cell types of different developmental origins including tangentially migrated prethalamic, preoptic, and hypothalamic ones (Moreno & Gonzalez, 2007; Aerts & Seuntjens, 2021; Medina et al., 2023). As a result, cell types within the different nuclei can substantially differ in their expression profile and connectivity. Most of the subpallial EA which consists of the medial amygdala (MeA), the central amygdala (CeA) and the bed nucleus of the stria terminalis (BST), originate from the lateral, medial and caudal ganglionic eminences (LGE, MGE and CGE) in tetrapods. When discussing the neuroanatomy of the teleostean amygdala, one has to consider that the concept of “the amygdala” has shifted over time and evolved with an increased understanding of the prosomeric organization of vertebrate forebrains. In addition, the telencephalon of teleosts differs substantially from mammals due to its outward growing process eversion and more basally derived molecular makeup (Wullimann & Mueller, 2004; Nieuwenhuys, 2009; Mueller et al., 2011; Folgueira et al., 2012; Furlan et al., 2017). Based on topology and molecular marker expression, a prosomeric ground plan of the zebrafish amygdala has been identified that defined a number of homologs to mammalian amygdaloid nuclei (Northcutt, 1995; Ganz et al., 2012; Porter & Mueller, 2020). These include regions corresponding to CeA and MeA amygdaloid nuclei, as well as the teleostean equivalent of the BST in the dorsal (Vd(d)), central (Vc), posterior (Vp) and supracommissural (Vs) divisions of the ventral telencephalon.

One controversially discussed and often overlooked territory represents the *isl1*-negative Vdd region described for carp-like (cyprinid) goldfish and zebrafish (Northcutt, 2006). Prior studies have considered the Vdd region in zebrafish and goldfish a dorsal striatal region (Northcutt, 1995; Ganz et al., 2012). However, the dorsal striatum (caudate and putamen) of mammals is defined as a derivative of the *Isle*t-1 positive ventral LGE (vLGE) and the majority of dorsal striatal neurons develops from *Islet-1* expressing precursor cells. We therefore postulated that the teleostean Vdd region represents an extended portion of the dorsal LGE (dLGE) that has not been identified or may differ from the one in tetrapods (Porter & Mueller, 2020). Correctly delineating the teleostean dLGE is crucial for recognizing the pallial-subpallial border (PSB) and the likely location of the anti-hem closely associated with the development of the rostral portion of the subpallial central amygdala topologically adjacent to the basolateral complex in mammals. Such definitions are also critical for understanding the morphogenetic processes underlying telencephalic eversion and ultimately will help understanding the developmental constraints of amygdala evolution in vertebrates.

To better characterize the subpallial DA system in zebrafish and the evolution of the amygdala, we analyzed the distribution of Th immunoreactive neurons in comparison to markers for distinct subpallial territories at larval stages. Our analyses of transcription factor expression locate these DA neurons to the dorsalmost compartment of the subpallium, directly adjacent to PSB. The DA neurons are located in a domain of *calb2a* expression, characteristic for the extended medial amygdala (EMeA). Indeed, a DA subpopulation expresses *calb2a,* demonstrating that the zebrafish subpallial DA system is located in the extended medial amygdala. Within the EMeA, a subpopulation of DA neurons expresses *sst7*, revealing diversity of this DA group. We present a detailed anatomical model for the EMeA DA neurons in the 5 days post fertilization (dpf) larval brain. Our findings provide a basis to investigate the development and function of the intratelencephalic DA system evolved in the subpallial amygdala of teleosts.

## 4. Material and Methods

### 4.1 Zebrafish strains, maintenance and specimen preparation

Wildtype zebrafish of the ABTL strain (www.ZFIN.org ID ZDB-GENO-031202-1) were kept under standard conditions (Westerfield, 2000). Embryos obtained by natural breeding were raised at 28.5 °C in 3 g/l Red Sea salt (Red Sea Aquatics Ltd) and staged according to Kimmel (Kimmel et al., 1995). To inhibit pigmentation, 0.2 mM n-phenylthiourea was added to the media. At 5 dpf, larvae were fixed in 4 % Paraformaldehyde (PFA) in PBST (PBS + 0.1 % Tween 20) at 4 °C overnight. After fixation, fish were washed in PBST and brains were dissected. Dissected brains were stored in 100 % methanol at -20 °C until needed. 30 dpf fish were treated with MS-222 (0.300 mg/ml) until cardiac arrest, followed by decapitation. Subsequently, the brains were dissected, fixed and stored as described for 5 dpf larvae. All experiments were carried out in accordance with the German Animal Welfare Act.

### 4.2 Cloning of cDNA fragments and *in-situ* probe synthesis

To extract RNA from 3 dpf embryos of the ABTL strain, the Qiagen RNeasy Mini Kit (Qiagen, ID: 74004) was used according to manufacturer’s instructions. cDNA was synthesized using the SuperScript™ III Reverse Transcriptase Kit (Invitrogen, 18080093) using oligo(DT)_20_ primer according to manufacturer’s instructions. Coding sequence fragments of *calb2a* (ENSDART00000060160.5), *lhx8a* (ENSDART00000019078.5)*, sst7* (ENSDART00000047378.7) and *tbr1b* (ENSDART00000006612.7) were amplified using MyTaq™ DNA Polymerase (Bioline, BIO-21107) and the following primers:

*calb2a* fwd: 5’-GAAATACGATACAGACCGCAGC-3’

*calb2a* rev: 5’-GCCATGATGCTTTGCTTGTAAC-3’

*lhx8* fwd: 5’-AGCCTGTCATACTGGACAATG-3’

*lhx8* rev: 5’-CATGCTGTCCTCTGACCTGA -3’

*sst7* fwd: 5’-CAGCACAAAGTAGGGAGTTGAG-3’

*sst7* rev: 5’-AGAGTAAGTCCACGGAGACC-3’

*tbr1b* fwd: 5’-TGGCTACCCAAACGCACAAG-3’

*tbr1b* rev: 5’-GCGGAGGAAAACTGGTAGAAAG-3’

Expected PCR product sizes were identified by agarose gel electrophoresis, cloned into the pCRII-TOPO plasmid (Invitrogen, K461020) and cDNA sequence verified by sequencing. Probes used in this study are given in Table S1.

### 4.3 Fluorescent whole mount *in-situ* hybridization (WISH)

WISH was performed as described previously with minor modifications (Filippi et al., 2007; Ronneberger et al., 2012). Brains were rehydrated in decreasing concentrations of methanol, washed with PBST and incubated with 1 % hydrogen peroxide in PBST for 30 minutes at room temperature (RT). After several PBST washing steps, permeabilization was performed by incubating in 4 % Triton™ X-100 (Applichem, A1388) in PBST for 60 minutes at RT. After rinsing with PBST, brains were post-fixed in 4 % PFA / PBST for 30 minutes and washed several times with PBST. For prehybridization, brains were incubated in hybridization buffer (50 % formamide, 5x SCC, 50 µg/ml heparin, 5 mg / ml torula RNA, 0.1 % Tween 20) at 65 °C for at least 4 hours. DIG- and DNP-labelled probes were diluted in hybridization buffer and hybridization was performed overnight at 65 °C. The following day, unbound probe was washed out using decreasing concentrations of formamide in SSC at 65 °C. After washing in TNT (100 mM Tris-HCl pH 7.5, 150 mM NaCl, 0.5 % Tween 20) at RT, brains were incubated with 1 % Blocking Reagent (Roche, #1096176) for 2 hours and incubated with 1:400 Anti-Digoxigenin-POD (Roche, #11207733910) overnight at 4 °C. Brains were then washed several times with TNT and stained for 45 minutes by a tyramide-Alexa 488 or 555 working solution according to manufacturer’s instructions (Invitrogen, B40953/B40955). After several TNT washes, we proceeded with Th-immunohistochemistry or continued with the staining of the DNP-probe as follows. The peroxidase of the Anti-Digoxigenin-POD antibody was inactivated by incubation in 1 % hydrogen peroxide in TNT for 30 minutes at RT. Brains where washed several times in TNT followed by blocking as described above and incubation in 1:100 Anti-DNP-HRP conjugate (Akoya Biosciences, TS-000400) overnight at 4 °C. On the following day, brains were washed in TNT followed by incubation in tyramide-Alexa 488 or 555 working solution for 90 minutes and washed several times in TNT. We proceeded with the Th-immunohistochemistry as described below.

### 4.4 Immunohistochemistry

Brains previously stained by fluorescent *in-situ* hybridization were equilibrated in PBTD (1 % DMSO in PBST) and incubated in antibody blocking solution (5 % goat serum (Sigma Aldrich, #G9023), 1 % Blocking Reagent (Roche, #1096176), 1 % BSA (Sigma Aldrich #A6003)) for 2 hours at RT. Brains were incubated overnight at 4 °C with 1:500 polyclonal rabbit-anti-Th primary antibody (Ryu et al., 2007) in blocking solution. On the following day, brains were washed several times with PBTD containing 200 mM KCl. After overnight incubation with 1:1000 secondary goat-anti-rabbit IgG Alexa 633 (Invitrogen, A21070) in antibody blocking solution, brains were washed in PBTD containing 200 mM KCl followed by several PBST washing steps and stored in 80 % Glycerol / PBST at 4 °C until confocal imaging.

### 4.5 Specimen clearing and hybridization chain reaction (HCR)

Dehydrated 30 dpf brains were rehydrated in decreasing concentrations of methanol in PBST and washed several times in PBST. Afterwards, brains were incubated for 45 minutes in DEEP-Clear Solution-1.1 (Pende et al., 2020) at 37 °C. After rinsing with PBST, brains were postfixed with 4 % PFA / PBST for 20 minutes at RT. After several PBST washing steps, multiplexed hybrid chain reaction (HCR) was performed according to the manufacturers protocol v3.0 (Molecular Instruments). HCR probes for *calb2a* (ENSDART00000060160.5) and *th* (ENSDART00000040410.7) detection are given in Supplementary Table S2.

### 4.6 Imaging and figure preparation

Samples for imaging were mounted in 80 % Glycerol / 20 % PBST /1.5% low melting Agarose (Biozym Scientific, 850080). Imaging was performed on the Zeiss LSM 880 confocal microscope (Life Imaging Center, University of Freiburg) using the LD LCI Plan-Apochromat 25x/0.8 NA Imm Corr DIC M27 objective and the Zeiss ZEN Black 2.3 software. Stacks were recorded with a resolution of 500×500 pixel with 0.66 µm x 0.66 µm per pixel. Stacks were either recorded with a z-plane thickness of 1 µm (Fig. 1a,b; 1e,f; 2a-c; 2f,g; 3a-d; 6a,b) or 2 µm (Fig. 1c,d; 2d,e; 4; 5; 6c,d; 7). Detector range for the recording of Alexa 488 was 499-553 nm, for Alexa 555 570-624 nm and for Alexa 633 642-686 nm. Typically, 2-3 brains for each analysis were documented (exact numbers for each analysis is provided in the respective figures). The scales were generated from the metadata of the ZEN image stacks using the Fiji distribution of ImageJ (Schindelin et al., 2012) and for visibility adjusted in their thickness using Adobe Illustrator CC. Brightness of images was linearly adjusted using Fiji. For better visualization the brightness of the blowups was further adjusted linearly. Transversal and sagittal views were generated from horizontal stacks with the computational reslicing plugin (Identifier: legacy:ij.plugin.Slicer) in Fiji. Figures were assembled and the schematic brain model was generated using Adobe Illustrator CC.

**Figure 1:**
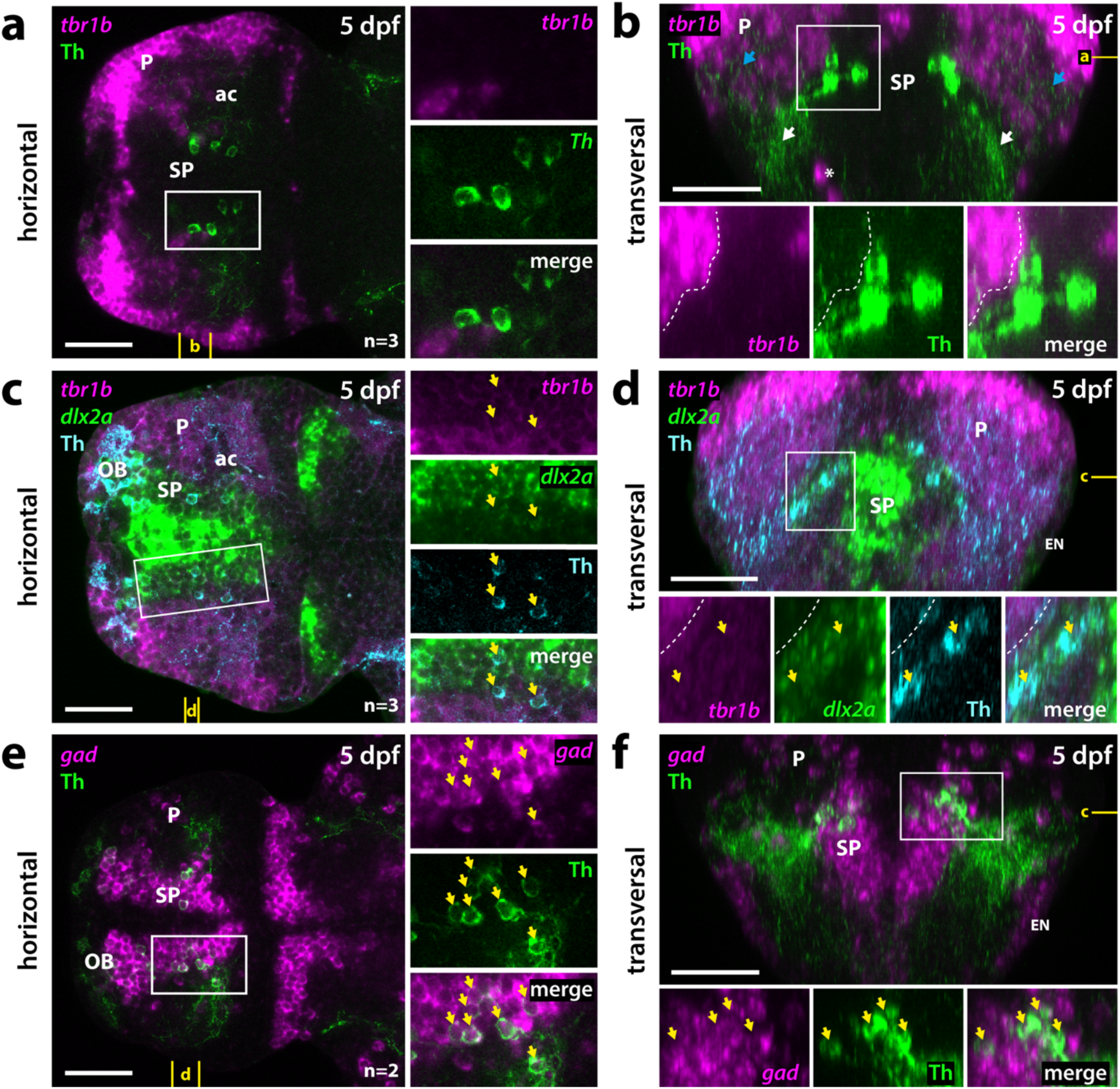
Validation of the subpallial localization of telencephalic DA neurons. (a-f) Fluorescent *in-situ* hybridization combined with anti-Th immunofluorescence in dissected 5 dpf zebrafish brains. **(a+b)** The pallial marker *tbr1b* **(a)** Horizontal plane at the position marked in yellow in (b). **(b)** Transversal projection as marked in yellow in (a). White asterisk marks scattered septal *tbr1b* positive cells. Blue arrows mark Th immunoreactive fibers in pallial areas. White arrows mark Th immunoreactive fibers in lateral parts of the ventral subpallium. Dotted line marks the border of the *tbr1b* expressing pallium. **(c+d)** Staining for *tbr1b* and, as marker for subpallial regions, *dlx2a* shows that subpallial DA neurons are located at the PSB in Vd. Yellow arrows mark cells that express *dlx2a* and are immunoreactive for anti-Th. **(c)** Horizontal plane superficial to the anterior commissure at the position marked in yellow in (d). **(d)** Transversal projection as marked in yellow in (c). Dotted line marks the border of the *tbr1b* expressing pallium. **(e+f)** Combined probes for *gad1a*, *gad1b* and *gad2* together mark all GABAergic neurons, and in combination with immunostaining for Th confirm the GABAergic cotransmitter type of subpallial DA neurons. Yellow arrows mark cells positive for *gad* and immunoreactive for anti-Th. **(e)** Horizontal plane superficial to the anterior commissure at the position marked in yellow in (f). **(f)** Transversal projection as marked in yellow in (e). **(a-f)** White boxes mark the areas presented in the blowups. n = number of analyzed brains per experiment. ac = anterior commissure, dpf = days post fertilization, EN = entopeduncular nucleus, OB= olfactory bulb, P = pallium, SP = subpallium. Scale bars = 50 µm.

**Figure 2:**
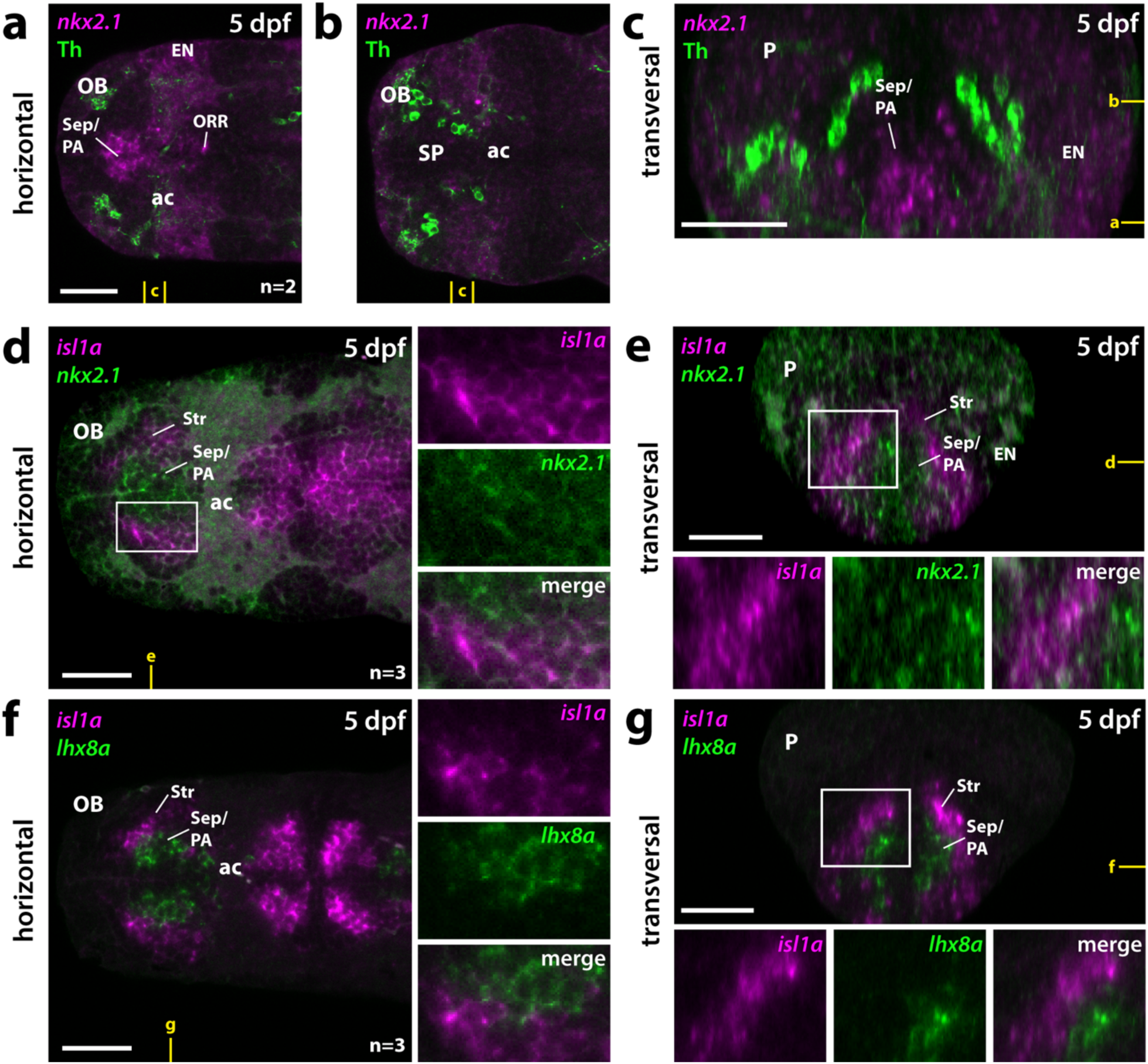
Subpallial DA neurons do not locate to MGE-derived septopallidal regions. (a-c) Fluorescent *in-situ* hybridization for *nkx2.1* that marks MGE derivatives in combination with Th immunostaining indicates that subpallial DA neurons are located dorsolateral of MGE derived septopallidal regions. **(a+b)** Horizontal plains at the z-positions depicted in yellow in (c). **(c)** Transversal projection as marked in yellow in (a+b). **(d+e)** Double staining for *nkx2.1* and *isl1a* marks MGE septopallidal derivatives and the vLGE-like striatum, respectively. **(d)** Horizontal plain at the z-position depicted in yellow in (e). **(e)** Transversal plane as marked in yellow in (d). **(f+g)** Double fluorescent *in-situ* hybridization for *isl1a* and *lhx8a* validates the distinct localization of MGE and vLGE derived regions. **(f)** Horizontal plain at the z-position depicted in yellow in (g). **(g)** Transversal plane as marked in yellow in (f). **(a-g)** White boxes mark the areas presented in the blowups. n = number of analyzed brains per experiment. ac = anterior commissure, dpf = days post fertilization, EN = entopeduncular nucleus, OB= olfactory bulb, ORR = optic recess region, P = pallium, PA = pallidum, Sep = septum, SP = subpallium, Str = striatum. Scale bars = 50 µm.

**Figure 3.**
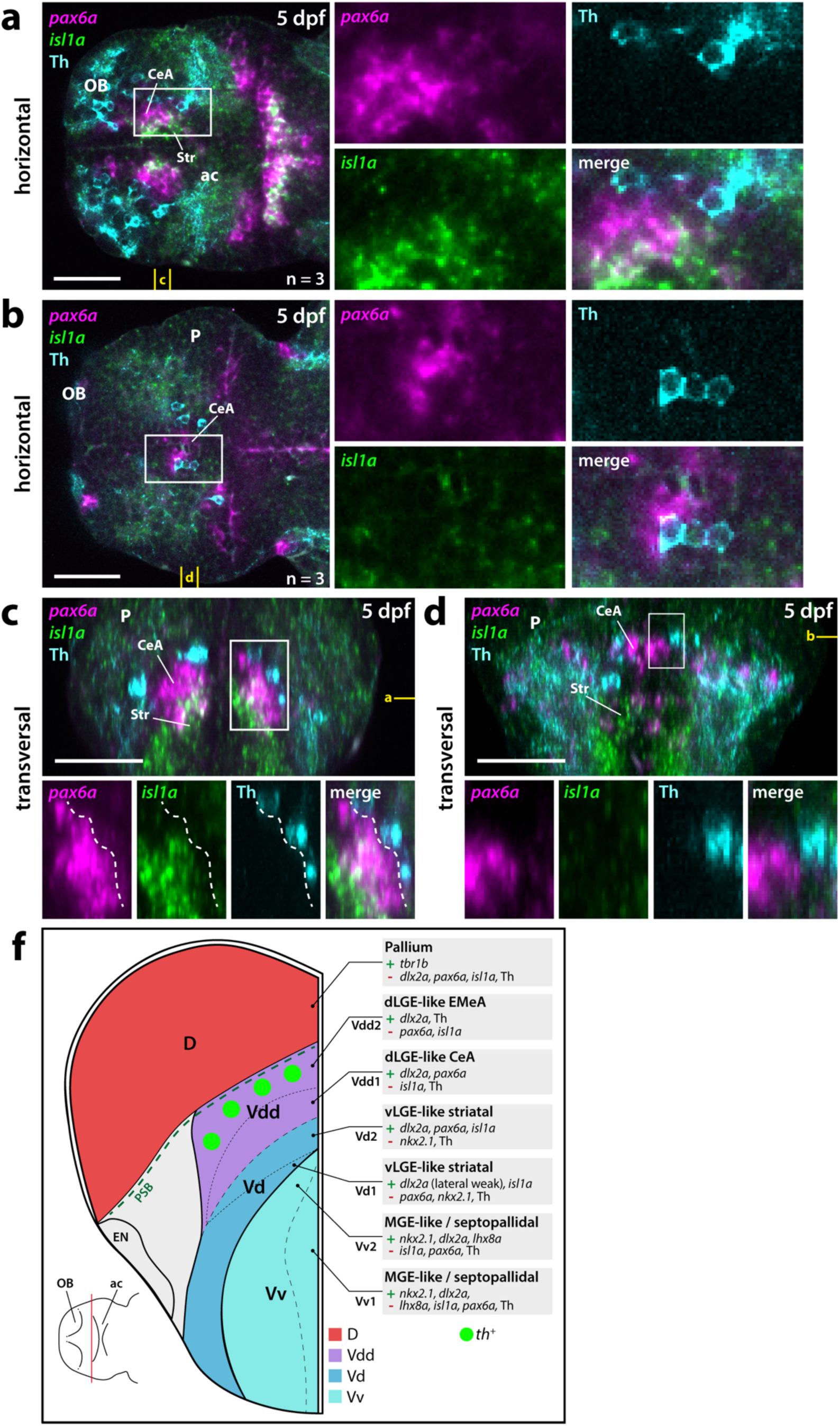
Subpallial DA neurons locate dorsolateral to both vLGE and the pax6 positive regions of dLGE. (a-d) Double fluorescent *in-situ* hybridization for *isl1a* and *pax6a* combined with Th immunofluorescence in 5 dpf zebrafish brains. See also Supplementary Video 1. **(a+b)** Horizontal planes at the positions marked in yellow in (c+d). **(c+d)** Transversal projections at the position marked in yellow in (a+b). Dotted line in (c) marks the border of the *pax6a* expressing Vdd. **(a-d)** White boxes mark the areas presented in the blowups. **(f)** Schematic transversal representation summarizing data from Figures 1-3. The subpallial dopaminergic neurons are located primarily dorsolateral of *isl1a* and *pax6a* positive derived striatal and CeA derivatives. n = number of analyzed brains per experiment. ac = anterior commissure, CeA = central amygdala, D = Pallium, dpf = days post fertilization, dLGE = dorsal lateral ganglionic eminence, EMeA = extended medial amygdala, EN = entopeduncular nucleus, MGE = medial ganglionic eminence, OB = olfactory bulb, P = pallium, PSB = pallial-subpallial border, Str = striatum, V = ventral telencephalic area, Vd = dorsal division of the ventral telencephalon, Vdd = dorsalmost division of the ventral telencephalon, Vv = ventral division of the ventral telencephalon. Scale bars = 50 µm.

**Figure 4:**
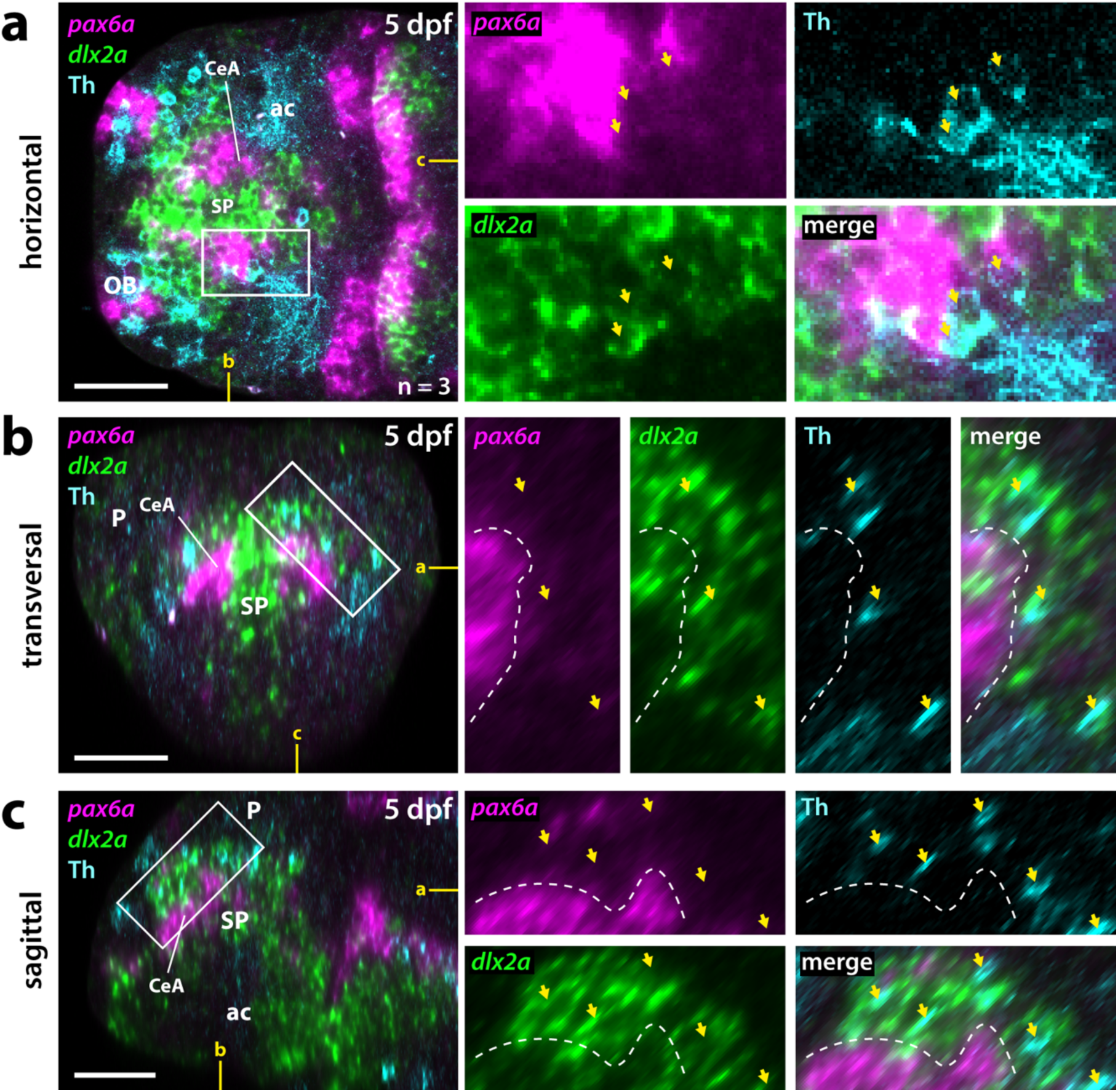
The subpallial DA neurons are located within an area of Vdd devoid of *pax6a* expression. (a-c) Double fluorescent *in-situ* hybridization for *pax6a* and *dlx2a* combined with Th immunofluorescence in 5 dpf brains. **(a)** Horizontal plane as depicted in yellow in (b+c). **(b)** Transversal plane as depicted in yellow in (a+c). **(c)** Sagittal plane as depicted in yellow in (a+b). **(a-c)** White boxes mark the areas presented in the blowups. Yellow arrows mark anti-Th immunoreactive cells. Dotted line in (b+c) marks the border between the *pax6a* expressing and *pax6a* free regions of Vdd. n = number of analyzed brains per experiment. ac = anterior commissure, CeA = central amygdala, dpf = days post fertilization, OB= olfactory bulb, P = pallium, SP = subpallium. Scale bars = 50 µm.

**Figure 5:**
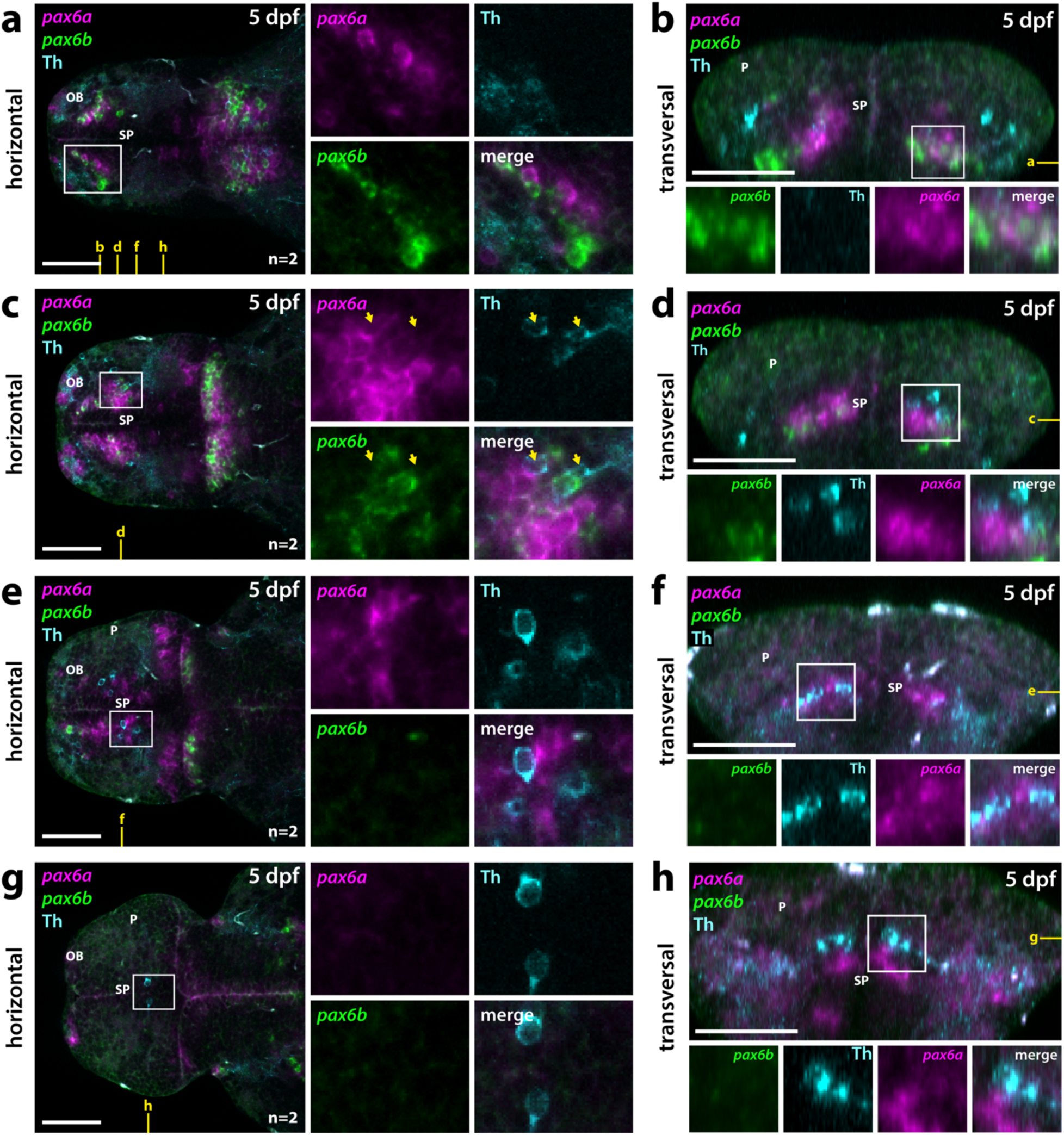
The pax6a/b are differentially expressed in the zebrafish telencephalon, but there is no co-expression with Th. (a-h) Double fluorescent *in-situ* hybridization for *pax6a* and *pax6b* combined with Th immunofluorescence in dissected brains. **a – c – e – g** show progressively more dorsal horizontal planes, **b – d – f – h** show progressively more caudal transversal sections (see yellow marks in **a;** corresponding horizontal and transversal images are indicated in each panel). **(a-b**) In the rostral subpallium, predominantly *pax6b is expressed.* **(c-d)** At more caudal positions there is an intermingled expression of the two *pax6* paralogs. Yellow arrows mark anti-Th immunoreactive neurons. **(e-h)** In the caudal subpallium and at supracommissural levels only *pax6a* expression can be detected. White boxes mark the areas presented in the blowups. n = number of analyzed brains per experiment. dpf = days post fertilization, OB= olfactory bulb, P = pallium, SP = subpallium. Scale bars = 50 µm.

**Figure 6:**
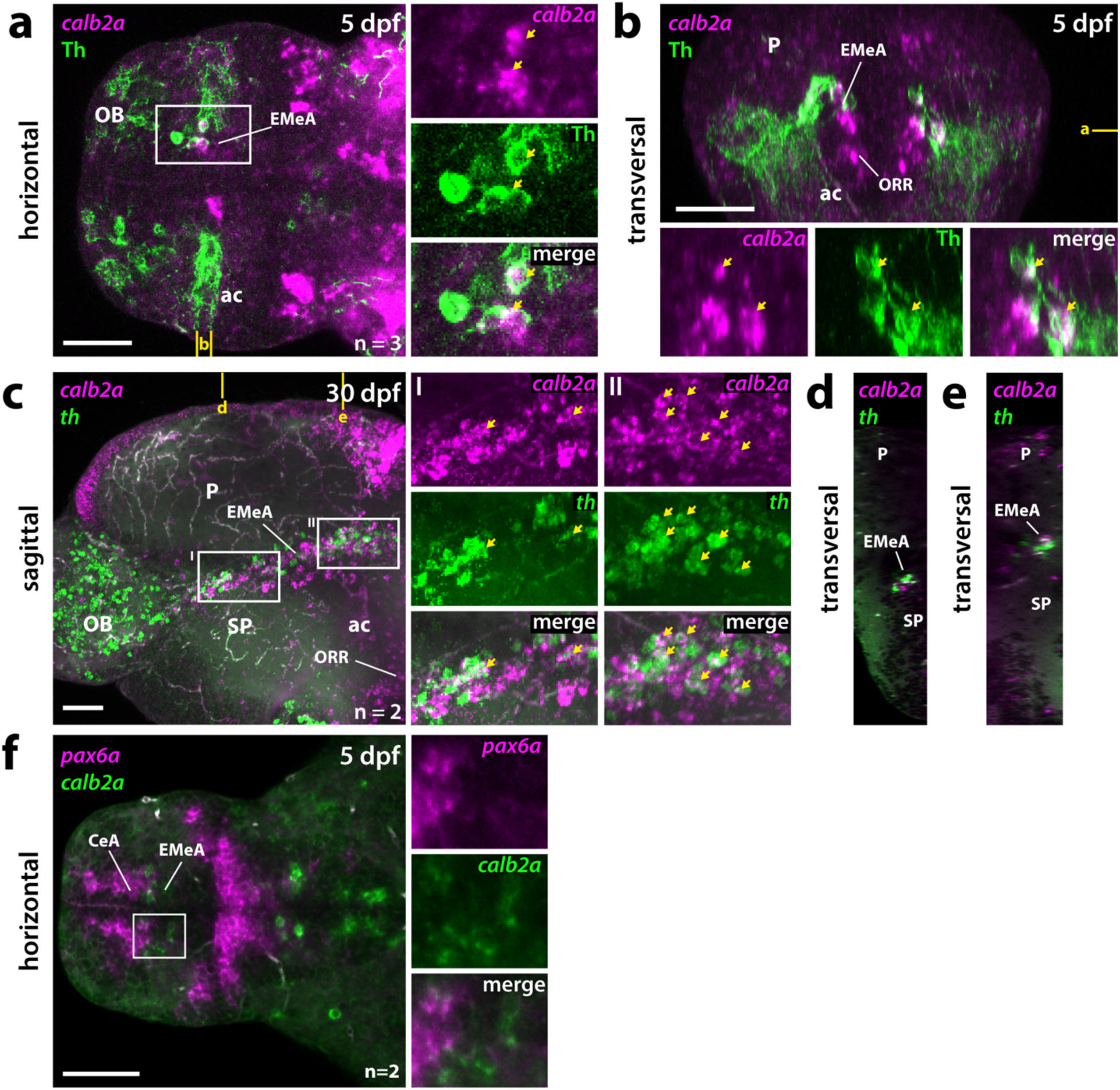
Calbindin expression localizes subpallial DA neurons to the medial extended amygdala. (a-e) WISH for *calb2a* in combination with Th immunofluorescence in 5 dpf (a,b) and 30 dpf (c-e) brains. **(a,b)** Subpallial dopaminergic neurons are located within a domain of the EMeA defined by *calb2a* expression. A subpopulation coexpresses *calb2a* and Th. See also, Supplementary Video 2. **(a)** Horizontal plain at the z-position depicted in yellow in (b). **(b)** Transversal projection as marked in yellow in (a). Coexpressing cells are marked by yellow arrows. **(c-e)** In the 30 dpf brain *calb2a* is coexpressed in many *th* expressing neurons and *calb2a* positive cells intermingle with DA neurons along the entire rostro-caudal axis of the EMeA. **(d+e)** Transversal planes at the positions depicted in (c). Coexpressing cells are marked by yellow arrows. **(f)** Double WISH for *calb2a* and *pax6a. calb2a* positive neurons do not coexpress *pax6a*. **(a-f)** White boxes mark the areas presented in the blowups. n = number of analyzed brains per experiment. ac = anterior commissure, CeA = central amygdala, dpf = days post fertilization, EMeA = medial extended amygdala, OB= olfactory bulb, ORR = optic recess region, P = pallium, SP = subpallium. Scale bars = 50 µm.

**Figure 7:**
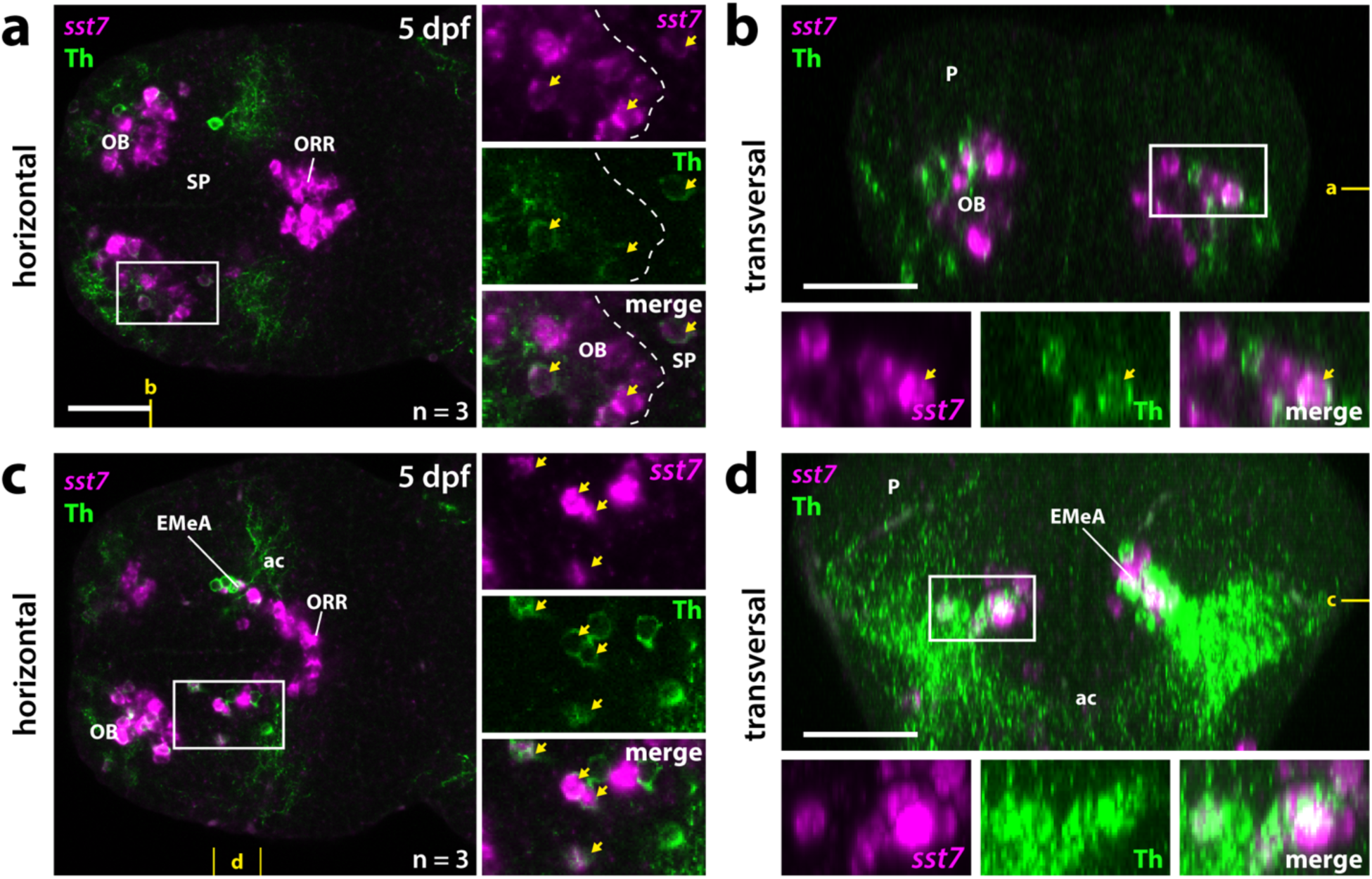
A subset of subpallial DA neurons express *somatostatin family member 7*. *(sst7)*. (a-d) WISH for *sst7* combined with anti-Th immunofluorescence in 5 dpf brains. **(a,b)** *sst7* is expressed within the olfactory bulb and optic recess region. A subset of olfactory bulb DA neurons coexpresses Th and *sst7.* See also Supplementary Video 3. **(a)** Horizontal plane at the z-position marked in yellow in (b). Dotted line separates the olfactory bulb *sst7* expressing cells from the one in the subpallium. **(b)** Transversal plane as marked in yellow in (a). (**c,d)** A subset of subpallial DA neurons coexpresses *sst7.* **(c)** Horizontal plain at the z-position marked in yellow in (d). **(d)** Transversal projection as marked in yellow in (c). **(a-d)** White boxes mark the areas presented in the blowups. Coexpressing cells are marked by yellow arrows. n = number of analyzed brains per experiment. ac = anterior commissure, EMeA = extended medial amygdala, OB= olfactory bulb, ORR = optic recess region, P = pallium, SP = subpallium. Scale bars = 50 µm.

## 5 Results

### 5.1 Subpallial dopaminergic neurons have a GABAergic cotransmitter type in 5 dpf zebrafish larvae

In 5 dpf zebrafish telencephalon, anti-Th immunoreactive DA neurons can be detected in the Vd(d) and Vp, directly adjacent to *tbr1b* expressing regions likely corresponding to the medial (Dm) and posterior division (Dp) of the pallium (Figure 1a+b). As previously reported, the subpallial DA neurons project locally, into lateral parts of the ventral subpallium, and into the anterior commissure (Figure 1b, white arrows) (Tay et al., 2011). However, substantial anti-Th immunoreactive innervation can also be detected in *tbr1b* positive pallial regions (Figure 1b, blue arrows). Scattered *tbr1b* expressing cells can be detected in the ventral subpallium (Vv), which might represent septal cells of pallial origins as described previously (Figure 1b, white asterisk) (Mione et al., 2001; Mueller et al., 2008; Filippi et al., 2012; Ganz et al., 2012) The subpallial DA neurons are positioned medially adjacent to *tbr1b* positive pallial regions, and thus locate to the most dorsolateral portion of the *dlx2a* positive Vd (Figure 1c,d). Dlx2 marks the ventral forebrain, including subpallial areas, throughout vertebrates (Puelles et al., 1999; Mueller et al., 2008; Moreno et al., 2009).

Subpallial DA neurons have a GABAergic cotransmitter-type (Filippi et al., 2014), which is in line with our observed coexpression of Th and *gad* (combinatorial staining of *gad1a*, *gad1b* and *gad2*) in the subpallium at 5 dpf (Figure 1e,f). The laterally displaced *dlx2a* and *gad2* positive areas likely correspond to the entopeduncular nucleus (EN) (Figure 1d,f,) (Solek et al., 2017; Wullimann, 2022). Taken together the subpallial DA neurons of 5 dpf zebrafish larvae are located in the dorsal subpallium medioventral adjacent to the *tbr1b* positive Dm and Dp and have a GABAergic cotransmitter type.

### 5.2 Subpallial dopaminergic neurons are located dorsolateral of *nkx2.1* and *lhx8a* positive MGE derivatives

To identify the exact localization and potential origin of the subpallial DA neurons, we performed coexpression studies of Th with transcription factors and differentiation markers that label distinct subpallial regions and lineages. We also analyzed the septopallidal region of Vv, in order to translate our observations into a model of the larval zebrafish subpallium at 5 dpf (Figure 3f).

In mice, the transcription factor Nkx2.1 specifies MGE and preoptic area derived telencephalic regions such as the pallidum, septum as well as the EN (Sussel et al., 1999; Puelles et al., 2000; Xu et al., 2008). Likewise, in the zebrafish telencephalon *nkx2.1* is expressed in the preoptic area as well as in pallidal and septal areas within Vv, including territories posterior to the anterior commissure recently termed as the optic recess region (ORR) (Ganz et al., 2012; Manoli & Driever, 2014; Affaticati et al., 2015). Subpallial DA neurons are neither located within *nkx2.1* positive regions, nor we can detect coexpression of Th with *nxk2.1* (Figure 2a+b) (Filippi et al., 2012). The subpallial DA neurons are clearly located dorsolateral to *nkx2.1* expressing areas (Figure 2c). These observations confirm that the subpallial DA neurons are not located in pallidal or septal regions and that it is likely that they are not part of the *nkx2.1* positive MGE / ORR lineage. Dorsolateral of the *nkx2.1* positive regions in the Vv *isl1a* expression can be detected (Figure 2d+e). The *isl1a* positive regions in ventral telencephalic area (V) likely correspond to striatopallidal structures (Stenman et al., 2003).

Another molecular marker specifying the MGE is Lhx8 (Grigoriou et al., 1998). In Nkx2.1 mutant mice, the former MGE zone is dorsalized and acquires LGE characteristics, including the loss of Lhx8 expression (Sussel et al., 1999). A similar dorsalization was observed in *nkx2.1* morphant zebrafish larvae (Manoli & Driever, 2014). Double staining of *isl1a* and *lhx8a* shows that MGE (*lhx8a* positive) and vLGE (*isl1a*) derived regions are clearly distinct with few coexpressing cells at the interface of both regions (Figure 2f+g). While *nkx2.1* expression can be detected already at and adjacent to the ventricle, *lhx8a* expression is absent from the ventricular zone, but present more laterally, potentially in differentiated neurons of Vv (see scheme in Figure 3f). Together, these data demonstrate that the subpallial DA neurons are not located in MGE derived septopallidal regions of Vv.

### 5.3 Subpallial dopaminergic neurons are located dorsolateral of *isl1a* / *pax6a* positive LGE derivatives

Next, we asked in which anatomical subdivision of the dorsal subpallium, Vd(d) and more caudally Vs, the subpallial DA neurons are located. These regions contain the zebrafish homologs of mammalian vLGE (*isl1a*-positive) and dLGE (*pax6-*positive/ *isl1a*-negative) derived areas including the proposed teleostean striatum and parts of the subpallial amygdala (Puelles et al., 2000; Tole et al., 2005; Bupesh et al., 2011; Ganz et al., 2012). Vdd of adult zebrafish has been described to be a largely *isl1a* free region containing the teleostean subpallial amygdala composed of the CeA, MeA and BST homologues (Porter & Mueller, 2020). In zebrafish, the main portion of telencephalic *pax6a* positive cells is located in Vdd at the PSB, likely corresponding to the dLGE derived Pax6a positive cells in mammals that contribute to the striatum and CeA (Wullimann & Rink, 2001; Bupesh et al., 2011). Thus, we suggest that also in zebrafish *pax6a* can serve as a marker for these dLGE derivatives and parts of the CeA.

We found that subpallial DA neurons are dorsolateral of *isl1a* positive vLGE derived striatal areas in 5 dpf zebrafish (Figure 3a+c+d, Supplementary Video 1). This dorsolateral positioning is more distinct rostral to the anterior commissure for the *isl1a* positive Vv and Vd, but still visible at supracommissural levels of Vs (Figure 3c+d). Therefore, the DA neurons are not located in the *isl1* positive Vd. We next asked whether we can use *pax6* expression as marker to locate the DA neurons in specific Vdd anatomical entities.

Zebrafish have two *pax6* paralogs – *pax6a* and *pax6b* (Nornes et al., 1998), thus, due to subfunctionalization, *pax6b* may be expressed differentially to *pax6a*. To exclude the possibility that subpallial DA neurons may reside in a potential more dorsal *pax6b* expression domain, we analyzed the expression of both *pax6* paralogs by WISH combined with Th immunofluorescence. We observed differential expression of both *pax6* paralogs in the 5 dpf telencephalon. While *pax6a* is strongly expressed in the olfactory bulb, *pax6b* is absent in this region (Figure 5b). In the subpallium, at more rostroventral positions near the olfactory bulb, *pax6b* is predominantly expressed, while more caudally *pax6a* and *pax6b* double positive cells are located (Figure 5a-d). At supracommissural levels however, only *pax6a* can be detected while *pax6b* expression is largely absent (Figure 5e+h).

The subpallial DA neurons locate very close and often directly adjacent to *pax6a* positive regions of Vdd (Figure 3b, Supplementary Video 1). At supracommissural levels individual DA neurons almost intermingle with *pax6a* positive cells. However, the majority of the anti-Th immunoreactive subpallial DA neurons are more dorsolateral to *pax6a*. This area represents a *pax6a* negative region of Vdd marked by the expression of *dlx2a* (Figure 4a-c). For *pax6b,* we could detect intermingled *pax6b* and anti-Th immunoreactive cells (Figure 5c), however, most of the anti-Th immunoreactive cells are dorsolateral to the *pax6b* expressing cells (Figure 5d). Similar to *pax6a*, *pax6b* is not expressed in subpallial DA neurons. Thus, the subpallial DA neurons are located more dorsolateral to *pax6a/b* expressing regions of Vdd, which we term Vdd1, indicating the presence of a *pax6a*-negative region of the dLGE-like Vdd, termed here Vdd2 (Figure 2f). Vdd2 is *dlx2a* positive (Figure 1c,d and 4) and is positioned directly adjacent to *tbr1b* positive pallial regions. Thus, the DA neuron containing region Vdd2 in the subpallium is located directly adjacent to the pallial-subpallial border.

### 5.5 *calb2a* expression in a subset of subpallial dopaminergic neurons reveals localization in the extended medial amygdala

The calcium binding protein Calretinin has been described to be expressed in several nuclei of the mammalian amygdala (Rowniak, 2017). Within the subpallial amygdala, strong expression was detected in the medial nucleus. Also in zebrafish, the Calretinin homologs *calb2a/b* are expressed in the subpallium including Vd, Vdd, Vs and Vp (Castro et al., 2006). The expression of *calb2a/b* visualized by anti-Calretinin immunoreactivity has also served as a marker for the extended medial amygdala (EMeA) comprising the MeA and the BST in Vdd (Porter & Mueller, 2020).

We found that subpallial DA neurons in 5 dpf zebrafish larvae are intermingled with a dorsal subpopulation of *calb2a* expressing cells in Vdd. Indeed, a subpopulation of DA cells coexpresses Th and *calb2a* as judged from WISH (Figure 6a-b, Supplementary Video 2). This observation could be extended to the 30 dpf brain using HCR RNA detection, validating that at least a large subset of subpallial DA neurons is located in the *calb2a* positive EMeA (Figure 6c). A subset of DA neurons coexpress *calb2a* along the entire rostrocaudal extend of the EMeA at 30 dpf (Fig. 6c-e). We were not able to detect coexpression of *calb2a* and *pax6a* in the larval subpallium (Figure 6f). Thus, the zebrafish subpallial DA neurons are located in the EMeA (Vdd2), a Vdd territory devoid of *pax6a* expression.

### 5.6 A subset of subpallial DA neurons expresses *sst7*

In mammals, the neuropeptide Cortistatin is predominantly expressed by GABAergic interneurons of the cortex and the hippocampal formation, modulating sleep, learning and cortical excitability (de Lecea et al., 1996; de Lecea et al., 1997a; Tallent et al., 2005). In addition, individual Cortistatin producing neurons have been identified in the olfactory bulb, in the basolateral amygdala, as well as in subcortical structures such as the striatum and medial amygdala (de Lecea et al., 1997b). So far, the telencephalic distribution of the zebrafish Cortistatin orthologue Sst7 *(somatostatin family member 7)* has not been described in detail (Tostivint et al., 2004). In contrast to mammals, we were not able to detect *sst7* expression in the pallium of zebrafish larvae (Figure 7a-d, Supplementary Video 3). Rather, *sst7* expression is detected in the olfactory bulb, the subpallium and, caudal to the anterior commissure, to the ORR. In the subpallium *sst7* expression is restricted to the region which we consider to be the EMeA/BSTa in 5 dpf zebrafish. In fact, a subset of subpallial DA neurons also expresses *sst7* (Figure 5a-d). Thus, some subpallial DA neurons in EMeA likely produce the neuropeptide Sst7.

## 6. Discussion

The absence of mesodiencephalic DA systems in zebrafish raises the question whether there are any DA systems that provide substantial DA input into striatum, limbic systems and pallium similar to mammals (Smeets & Gonzalez, 2000; Yamamoto & Vernier, 2011). A potential source for DA is the subpallial DA group in zebrafish (Holzschuh et al., 2001; Kaslin & Panula, 2001; Filippi et al., 2010; Yamamoto et al., 2010), however, exact anatomical location, development and function of these neurons have not been well understood to this day. Here, we aim to identify the subpallial subdivisions in which the DA neurons reside, and define new markers to better resolve their identity. We focus our analysis on the 5 dpf early larval brain, when all major telencephalic subdivisions have established (Mueller & Wullimann, 2016), and the brain is still small enough such that the telencephalon can be documented entirely in a single confocal stack. We also analyze 30 dpf brains when the adult brain morphology has been established. Combined analyses of Th immunolocalization for DA neurons, *lhx8, dlx2a, nkx2.1, isl1a, pax6a,* and *tbr1b* as anatomical markers, as well as of *gad1a/1b/2, calb2a* and *sst7* as differentiation markers demonstrate that the zebrafish subpallial DA neurons are located in the extended medial amygdala, and with respect to *calb2a* and *sst7* constitute a diverse group of neurons.

### 6.1 Subdivisions of the larval zebrafish subpallium at 5 dpf

Previous studies on the subpallial topology of adult zebrafish defined homologies to MGE and vLGE/dLGE derived structures such as pallidal, striatal, and amygdaloid nuclei (Porter & Mueller, 2020). To understand their development, a more profound knowledge about the telencephalic anatomy at early stages is needed. Our analyses of *dlx2a, isl1a, lhx8, nkx2.1, pax6a,* and *tbr1b* expression in the telencephalon validate existing and suggest additional subdivisions, such that we propose six subdivisions of the ventral telencephalon rostral to the anterior commissure (Figure 8): two *nkx2.1* positive MGE-like septopallidal domains Vv, distinguished by expression of *lhx8a* in Vv2 only; two ventral LGE-like subdomains, a striatal Vd1 with *isl1* expression and a Vd2 with both *isl1* and *pax6* expression; and two dorsal LGE subdivisions Vdd, including the more ventral Vdd1 of central amygdala character with *dlx2a* and *pax6a* expression, but no *isl1,* and the Vdd2 of EMeA character which is *pax6a* negative but contains *calb2a* and sst7 positive cells.

**Figure 8:**
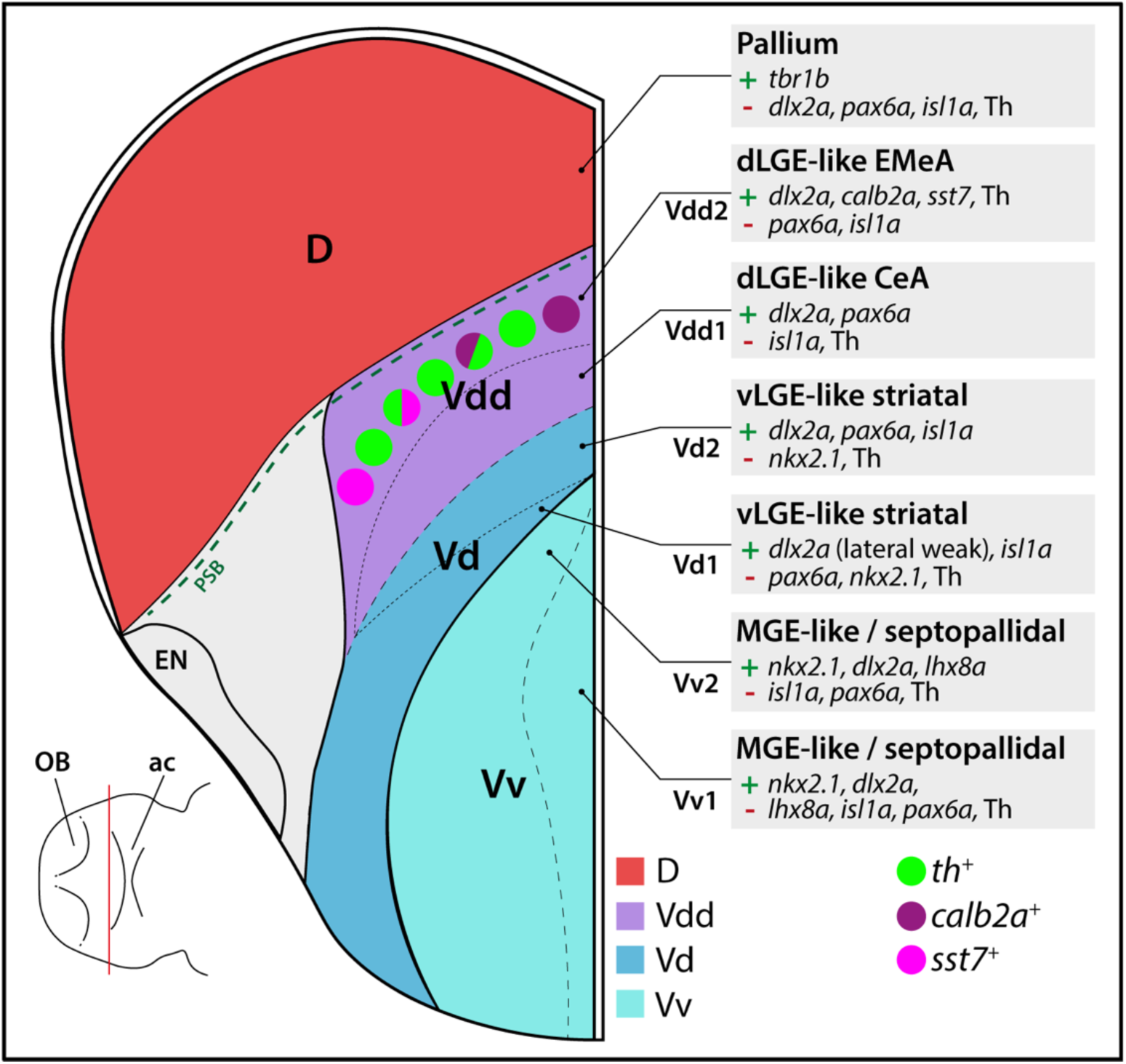
Schematic representation of localization of DA neurons relative to telencephalic divisions analyzed in this study. DA neuron subpopulations expressing calb2a or sst7 are indicated. Transversal repression of one hemisphere of the 5 dpf telencephalon. ac = anterior commissure, CeA = central amygdala, D = dorsal telencephalic area, dLGE = dorsal lateral ganglionic eminence, EMeA = extended medial amygdala, EN = entopeduncular nucleus, MGE = medial ganglionic eminence, OB = olfactory bulb, PSB = pallial-subpallial border, V = ventral telencephalic area, Vd = dorsal division of the ventral telencephalon, Vdd = dorsalmost division of the ventral telencephalon, Vv = ventral division of the ventral telencephalon, vLGE = ventral lateral ganglionic eminence.

These markers have been shown to identify ventral telencephalic subdivisions throughout vertebrate evolution. The knockout of the transcription factor Nkx2.1 in mice results in loss of MGE and an expansion of LGE derived tissues (Sussel et al., 1999), a highly conserved function (Puelles et al., 2000; Moreno, 2007). *Lhx8* is required for the specification of cholinergic cell types of the MGE lineage that reside within the pallidum (Zhao et al., 2003). The more dorsolateral located *isl1a* positive and *pax6a/lhx8a* negative region resembles the *isl1a* positive vLGE which gives rise to GABAergic projection neurons of the teleostean striatum and which was already described for zebrafish and many other vertebrates including rodents, amphibians and birds (Stenman et al., 2003; Moreno et al., 2008; Abellán & Medina, 2009; Ganz et al., 2012). The dorsalmost part of the zebrafish subpallium Vdd was described as *isl1a* free (Ganz et al., 2012; Porter & Mueller, 2020). The *isl1a* free Vdd expresses the transcription factor *pax6a*, likely resembling the Pax6 positive dLGE whose derivatives constitute parts of the central amygdala in tetrapods and also form part of the bed nucleus of the stria terminalis (BST) in mice (Yun et al., 2001; Abellán & Medina, 2009; Moreno et al., 2010; Bupesh et al., 2011). Thus, we suggest that the *pax6a* positive domain partially contributes to developing central amygdala-like structures of larval zebrafish. This is in line with the description of the subpallial amygdala of adult zebrafish as a largely *isl1a*-free GABAergic region ventral to the pallium (Porter & Mueller, 2020). Dorsolateral to the *pax6a* expression domain we could detect scattered cells expressing *calb2a*, which were described to form part of the teleostean equivalents of the EMeA in the adult brain (Porter & Mueller, 2020). In the same region we could observe numerous *sst7* neurons.

Thus, the subpallium of 5 dpf larval zebrafish already exhibits striking similarities to the subpallium of adult zebrafish with a high degree of regional identity and conservation of subpallial subdivisions of tetrapods. From an evolutionary perspective, our results support the notion that a tetrapod-like prosomerically organized amygdaloid complex originated in early vertebrates in a common ancestor of ray-finned fish and tetrapods (Porter & Mueller, 2020; Wullimann, 2022). The largely conserved topology of the zebrafish amygdala, as exemplified in this study for a portion of the subpallial EA, is the result of genetic and ontogenetic constraints shared between ray-finned fish and tetrapods.

### 6.2 Teleostean subpallial DA neurons are located within the extended medial amygdala

In contrast to mammals, zebrafish do not possess DA neurons in the midbrain but develop numerous *th* positive DA neurons in the subpallium (Guo et al., 1999; Holzschuh et al., 2001; Rink & Wullimann, 2002; Filippi et al., 2010). Considering that these subpallial DA neurons strongly innervate the zebrafish telencephalon, it is intriguing to investigate if zebrafish subpallial DA neurons have similar neuromodulatory functions as midbrain DA neurons of mammals (Tay et al., 2011). A recent study suggested the location of subpallial DA activity to the extended medial amygdala in the adult brain (Porter & Mueller, 2020), opening the exciting perspective that a portion of the zebrafish subpallial DA groups may be an evolutionary substitute of the mesolimbic system.

To test the hypothesis of the amygdaloid localization of the subpallial DA neurons, we analyzed the expression of marker genes suitable for defining subpallial regions in combination with distribution of Th immunoreactivity. The GABAergic subpallial DA neurons in 5 dpf zebrafish reside at the PSB medioventrally adjacent to *tbr1b* positive pallial regions. This *dlx2a* positive region of Vdd, Vp and Vs was described as the teleostean subpallial amygdala in adult zebrafish (Mueller et al., 2008; Ganz et al., 2012; Porter & Mueller, 2020). Porter and Mueller (2020) locate subpallial DA neurons to the adult EMeA based on their position dorsolateral to the *isl1a* positive Vd, which corresponds to vLGE derived striatal regions of mammals. Our finding that already in the larval brain the subpallial DA neurons are located dorsolateral to the *isl1a* expression domain corroborates their amygdaloid nature (Vdd). Interestingly, our data point to a novel early subdivision in Vdd. Previously, the *pax6a* expression domain was considered to form the PSB in zebrafish (Wullimann & Rink, 2002). Our findings however, suggest that a *dlx2a* positive and *pax6a* negative region of the dLGE-like Vdd forms at the PSB that we term Vdd2. We locate the subpallial DA neurons exactly to this *pax6a* and *isl1* negative Vdd2 area.

The Vdd2 area to which the subpallial DA neurons are located is characterized by dispersed *calb2a* (Calretinin homologue) expressing neurons, and indeed some of the DA neurons coexpress *calb2a.* In mammals Calbindin- and Calretinin expressing neurons populate major parts of various amygdaloid nuclei – including the EMeA (Wójcik et al., 2013; Rowniak, 2017). Likewise in adult zebrafish anti-Calretinin immunoreactivity marks the extended medial amygdala including the bed nucleus of the stria terminalis (Porter & Mueller, 2020). Thus, the presence of Vdd-derived *calb2*-expressing GABAergic neurons indicate the extended medial amygdaloid nature of these territories (EMeA/BSTa). Amongst others, the MeA is implicated in mediating fear responses as well as in the regulation of social behavior by receiving olfactory and vomeronasal inputs (Meredith et al., 1999; Herdade et al., 2006).

A further indication of subtype diversity in the EMeA DA population is provided by the expression of *sst7*. *sst7* codes for the somatostatin family member Sst7, an orthologue to the mammalian neuropeptide Cortistatin. Cortistatin expression can be detected in GABAergic interneurons of the cortex as well as in the hippocampus and in neurons of the olfactory bulb, the basolateral amygdala, the striatum, the BST, and the MeA (de Lecea et al., 1996; de Lecea et al., 1997a; de Lecea et al., 1997b; Tallent et al., 2005). Similarly, in the marsh frog *Rana ridibunda* the suggested Cort orthologue PSS2 is expressed in the medial pallium, but also in the preoptic area and the MeA (Tostivint et al., 1996). Our finding of *sst7* expression in 5 dpf zebrafish larvae EMeA and DA neurons indicates strong conservation.

### 6.3 Conclusions and further perspective

Our study advances two major aspects of subpallial organization and function. First, we refine anatomical organization of the dorsal LGE-like Vdd into a Vdd2 subdivision that may correspond to the EMeA and forms the PSB region, and a more ventral Vdd1 that is *pax6a* positive and may correspond to the CeA. Second, we locate the subpallial DA neurons to the EMeA both in larvae and mature zebrafish brains, postulating a potential “endolimbic” DA system in zebrafish. Mice embryos also have been shown to develop TH-immunoreactive neurons in the central and medial amygdala, only few of which persist into adulthood (Bupesh et al., 2013), indicating an early evolutionary origin of amygdaloid DA neurons. Our findings shed light on potential mechanisms of DA systems evolution. From an ancient progenitor that contained subpallial and mesdiencephalic DA systems, mammals evolved the more complex mesolimbic, mesostriatal and mesocortical systems that serve the needs of neuromodulation of complex behaviors. In contrast, teleost neuromodulation of behavior may have been driven more prominently by specific environmental stimuli rather than complex decision-making. In this line, there is a close link between olfactory control and behavior, the EMeA and its DA system. This may extend to social and other cues that ultimately provide input into the limbic system and in return are affected by limbic output.

The zebrafish subpallial DA neurons may also provide an interesting model to investigate local DA differentiation and establishment of connectivity within the subpallium. Derivation of DA progenitors from induced pluripotent stem cells for regenerative therapies of Parkinson’s disease has become a highly efficient procedure (Lee et al., 2000; Kim et al., 2021). Transplanted progenitors have the potential for long-term survival in the striatum (Li et al., 2016), and preclinical studies are ongoing (Piao et al., 2021).

## 7. Acknowledgements

We thank Dr. Eva Carl for designing and cloning the *lhx8a* cDNA containing *pCR2-TOPO* plasmid that we used for the generation of the *lhx8a* DIG probe used in this work. We thank Sabine Götter for excellent fish care and the Life Imaging Center and Roland Nitschke for assistance in confocal microscopy. Funded by the Deutsche Forschungsgemeinschaft (DFG, German Research Foundation) under Germany’s Excellence Strategy CIBSS—EXC-2189— Project ID: 390939984, and by DFG—322977937/GRK2344 MeInBio. Thomas Mueller was funded by the Human Frontiers Science Foundation Grant RGP0016/2019.

## Anatomical abbreviations

ac: anterior commissure
BST: bed nucleus of the stria terminalis
BSTa: anterior part of BST (in zebrafish)
CA: catecholaminergic, catecholamine
CeA: central amygdala
CGE: caudal ganglionic eminences
CNS: central nervous system
D: Dorsal telencephalic area (pallium)
Dm: dorsomedial divisions of the dorsal telencephalon
Dp: posterior division of the dorsal telencephalon
dStr: dorsal striatum
EN: entopeduncular nucleus
EMeA: extended medial amygdala
(v/d)LGE: (ventral/dorsal) lateral ganglionic eminences
MeA: medial amygdala
MGE: medial ganglionic eminences
OB: olfactory bulb
ORR: optic recess region
P: pallium
PA: pallidum
PSB: pallial-subpallial border
Sep: septum
SP: subpallium
V: ventral telencephalic area (subpallium)
Vc: central division of the ventral telencephalon
Vd: dorsal division of the ventral telencephalon
Vdd: dorsalmost division of the ventral telencephalon
Vp: posterior division of the ventral telencephalon
Vs: supracommissural division of the ventral telencephalon
vStr: ventral striatum
VTA: ventral tegmental area
Vv: ventral division of the ventral telencephalon

## Supplementary Materials

## Supplementary Tables

**Table S1:** Probes for WISH used in this study (page 2)

**Table S2:** HCR Probes used in this study (page 3)

## Supplementary Videos S1, S2 and S3

**Supplementary Video 1** (related to Figure 3) Subpallial DA neurons locate dorsolateral to both vLGE and the pax6 positive regions of dLGE. Double fluorescent *in-situ* hybridization for *isl1a* and *pax6a* combined with Th immunofluorescence in 5 dpf zebrafish brains. (Format MP4 1920 x 1080 pixel, 20.2 Mb).

**Supplementary Video 2** (related to Figure 6) Calbindin expression localizes subpallial DA neurons to the medial extended amygdala. WISH for *calb2a* in combination with Th immunofluorescence in 5 dpf brains. (Subpallial dopaminergic neurons are located within a domain of the EMeA defined by *calb2a* expression. A subpopulation coexpresses *calb2a* and Th. (Format MP4 1920 x 1080 pixel, 14.5 Mb).

**Supplementary Video 3** (related to Figure 7) A subset of subpallial DA neurons express *somatostatin family member 7 (sst7)*. WISH for *sst7* combined with anti-Th immunofluorescence in 5 dpf brains. *sst7* is expressed within the olfactory bulb and optic recess region. A subset of olfactory bulb DA neurons coexpresses Th and *sst7.* (Format MP4 1920 x 1080 pixel, 12.7 Mb).

---

**Table S3:**
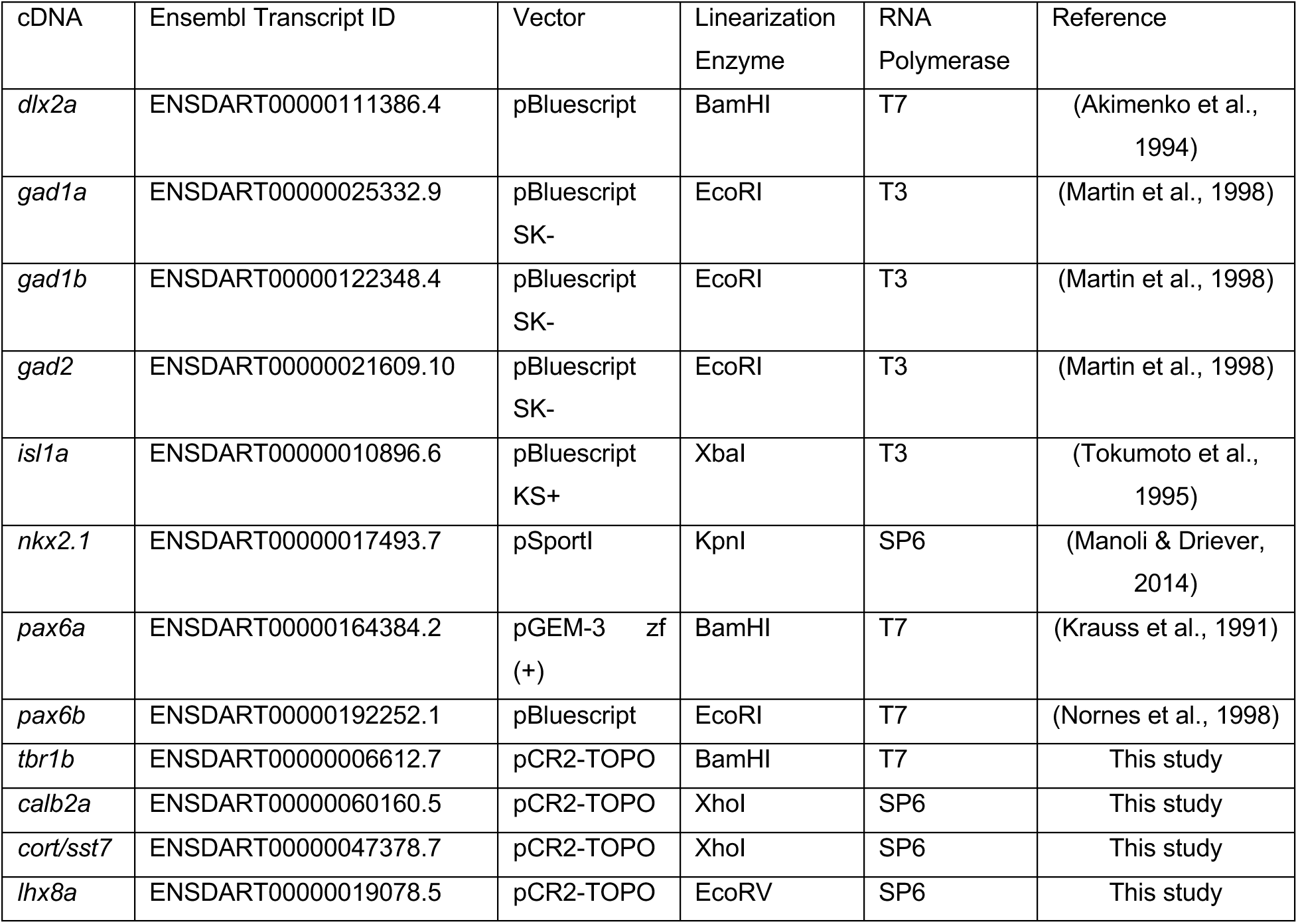
Probes for WISH used in this study.

| cDNA | Ensembl Transcript ID | Vector | Linearization Enzyme | RNA Polymerase | Reference |
| --- | --- | --- | --- | --- | --- |
| *dlx2a* | ENSDART00000111386.4 | pBluescript | BamHI | T7 | (Akimenko et al., 1994) |
| *gad1a* | ENSDART00000025332.9 | pBluescript SK- | EcoRI | T3 | (Martin et al., 1998) |
| *gad1b* | ENSDART00000122348.4 | pBluescript SK- | EcoRI | T3 | (Martin et al., 1998) |
| *gad2* | ENSDART00000021609.10 | pBluescript SK- | EcoRI | T3 | (Martin et al., 1998) |
| *isl1a* | ENSDART00000010896.6 | pBluescript KS+ | XbaI | T3 | (Tokumoto et al., 1995) |
| *nkx2.1* | ENSDART00000017493.7 | pSport1 | KpnI | SP6 | (Manoli & Driever, 2014) |
| *pax6a* | ENSDART00000164384.2 | pGEM-3 zf (+) | BamHI | T7 | (Krauss et al., 1991) |
| *pax6b* | ENSDART00000192252.1 | pBluescript | EcoRI | T7 | (Nornes et al., 1998) |
| *tbr1b* | ENSDART00000006612.7 | pCR2-TOPO | BamHI | T7 | This study |
| *calb2a* | ENSDART00000060160.5 | pCR2-TOPO | XhoI | SP6 | This study |
| *cort/sst7* | ENSDART00000047378.7 | pCR2-TOPO | XhoI | SP6 | This study |
| *lhx8a* | ENSDART00000019078.5 | pCR2-TOPO | EcoRV | SP6 | This study |

**Table S4:**
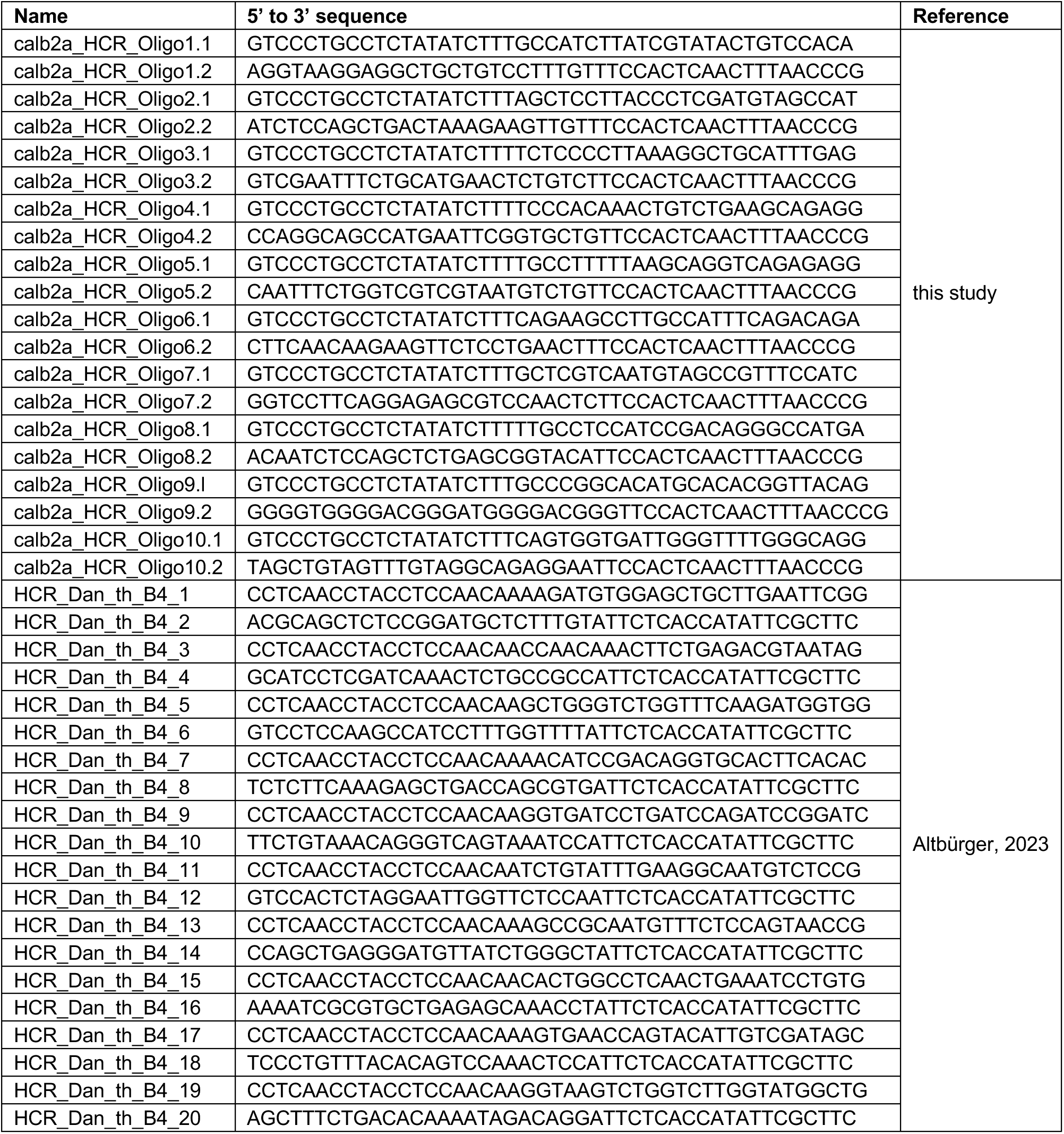
HCR Probes used in this study.

## Notes

### Competing Interest Statement

The authors have declared no competing interest.

